# NMR spectroscopy analysis reveals an altered metabolic homeostasis in Arabidopsis seedlings treated with a cytokinesis inhibitor

**DOI:** 10.1101/2020.06.21.163725

**Authors:** Thomas E. Wilkop, Minmin Wang, Angelo Heringer, Florence Zakharov, Viswanathan V. Krishnan, Georgia Drakakaki

**Affiliations:** Light Microscopy Core/Department of Physiology, University of Kentucky, Lexington, KY 40536; Department of Plant Sciences, University of California, Davis, CA 95616; Department of Chemistry, California State University, Fresno CA 93750; Department of Medical Pathology and Laboratory Medicine, University of California School of Medicine, Sacramento CA 95817

**Keywords:** cell division, plant primary metabolism, *Arabidopsis thaliana*, NMR-spectroscopy, cytokinesis inhibition

## Abstract

In plant cytokinesis, *de novo* formation of a cell plate evolving into the new cell wall partitions the cytoplasm of the dividing cell. Cell plate formation involves highly orchestrated vesicle accumulation, fusion, and membrane network maturation supported by the temporary integration of elastic and pliable callose. The small molecule, Endosidin 7 (ES7) arrests late cytokinesis in Arabidopsis by inhibiting callose deposition at the cell plate. Its effect is specific, as it does not broadly affect endomembrane trafficking or cytoskeletal organization. It has emerged as a very valuable tool for dissecting this essential plant process. In order to gain deeper insights regarding its mode of action and the effects of cytokinesis inhibition on overall plant growth, we investigated the effect of ES7 through a nuclear magnetic resonance spectroscopy metabolomics approach. In this case study, profiles of Arabidopsis leaf and root tissues were analyzed at different growth stages and ES7 exposure levels. The results show tissue-specific changes in the plant metabolic profile across a developmental gradient, and the effect that ES7 treatment has on the corresponding metabolome. The ES7 induced profile suggests metabolic compensations in central metabolism pathways in response to cytokinesis inhibition. Further, this study shows that long-term treatment of ES7 disrupts the homeostasis of primary metabolism in Arabidopsis seedlings, likely via alteration of hormonal regulation.

## Introduction

In a large-scale chemical genetics screening of small molecules interfering with endomembrane trafficking and polysaccharide deposition in Arabidopsis (Drakakaki et al., 2011), a number of highly specific compounds were identified. Among these compounds, Endosidin 7 (ES7), a heterocyclic organic molecule with attributes of both flavonoid and alkaloid derivatives, specifically inhibits callose deposition at the division plane and results in late-stage cytokinesis arrest (Park et al., 2014). Cytokinesis is a fundamental process of all life on earth, essential for plant growth and development. It is a highly regulated process, that involves the accumulation of membrane material coordinated with biopolymer deposition, such as callose, to structurally stabilize the maturing cell plate while it transitions into a new cell wall (Lipka et al., 2015; Samuels et al., 1995; Smertenko et al., 2017). Currently, little is known regarding the integration and coordination of polysaccharide deposition in conjunction with membrane maturation during cell plate expansion (McMichael and Bednarek, 2013; Drakakaki, 2015). Understanding the adjustment of the plant’s metabolome in response to ES7 treatment can provide insights into major metabolic fluxes during plant cytokinesis and potential compensating mechanisms.

The timely inhibition of cytokinesis by ES7 has been used to reveal the interplay of specific vesicle populations during cell plate assembly (Park et al., 2014; Davis et al., 2016). Using ES7 as a tool, the timely pattern of vesicle contributions during cell plate expansion can be dissected in order to better understand their role. This includes distinguishing between the early arrival of GTPase labeled cytokinetic vesicles and the vesicle fusion mechanisms visualized by the cytokinesis-specific SNARE protein, KNOLLE (Park et al., 2014). Furthermore, elevated levels of clathrin-coated vesicles accompany callose deposition during cytokinesis. The amount of these vesicles is reduced under ES7 treatment, suggesting they are stabilized through the temporal integration of callose (Park et al., 2014). The consistent effects of ES7 across the plant kingdom, from early diverging algae, e.g. Charophyte *Penium margaritaceum*, to higher plants, demonstrate that the pathways affected by ES7 are evolutionarily conserved (Davis et al., 2020). Previous research in Arabidopsis showed that the application of ES7 indirectly inhibits, callose synthase activity, namely the incorporation of UDP-glucose into β-1,3-glucan (Park et al., 2014).

*Arabidopsis thaliana* is a well-established model organism, employed in many studies aimed at understanding the biological functions across the plant kingdom (Meinke et al., 1998). A large number of detailed omics studies have been carried out to map and investigate the transcriptome, proteome, as well as metabolome of Arabidopsis during growth and development (Van Norman and Benfey, 2009; Joyce and Palsson, 2006; Hennig, 2007). This combined wealth of knowledge for Arabidopsis makes this model plant an excellent choice for performing untargeted metabolite analysis during plant development. The popularity of using Arabidopsis as a model system has led to several nuclear magnetic resonance (NMR)-based metabolomics studies in both solution and solid-state samples (Sekiyama et al., 2010; Kim et al., 2007; Gromova and Roby, 2010; Yuan et al., 2016). Approaches of ^1^H high-resolution magic angle spinning NMR, circumventing the need for elaborate sample preparation and allowing utilization of solid-state NMR spectroscopy for the study of intact leaves, were more recently developed (Augustijn et al., 2016, 2018). NMR-based methods for studying plant metabolomics, including sample preparation protocols and data analysis approaches, have been continuously refined (Kim et al., 2010; Deborde et al., 2019).

In order to understand the effect of ES7 on the overall plant development, physiology, and metabolism, we performed an NMR-based metabolomics analysis. Given the metabolism differences that exist between aerial tissues and roots, including photosynthetic activity, carbon assimilation, and nutrient acquisition (Thomas and Rodriguez, 1994; Koch, 1996; Abramoff and Finzi, 2015), we investigated both roots and leaves separately in our study. Metabolites were monitored in roots and leaves of treated and non-treated Arabidopsis seedlings, over a period of 4-10 days. In order to investigate the factors affecting metabolite levels across developmental stages and ES7 treatment, we utilized a partial least squares discriminant analysis (PLS-DA) for classification of the metabolites and a comprehensive multivariate statistical analysis to identify and quantify the metabolites that are differentially altered due to ES7 treatment. We found that the concentrations of over 50 metabolites are affected in Arabidopsis roots and leaves as a result of ES7 induced cytokinesis inhibition. Additionally, our work provides an NMR-based metabolomics protocol to study the effect of small molecules on plant metabolism.

## Material and Methods

### Plant materials and metabolite extraction

Arabidopsis seedlings were germinated on agar media with half-strength MS basal salts and 1% sucrose. Square plates were oriented vertically during growth. Seedlings were grown at 22°C with a 16h light cycle at ∼80 μE light intensity. ES7 was dissolved in DMSO and supplemented into the medium at concentrations of 0, 3, 5, and 10 μM.

During sample harvesting roots and leaves were separated by cutting with razor blades, and metabolites were extracted following a standard procedure (Orr et al., 2014; Kim et al., 2010). All seedlings in one plate contributed to an individual biological replicate. Briefly, tissue was homogenized with mortar and pestle in liquid nitrogen, and metabolites were extracted in methanol as follows. Homogenized tissue was lysed using 3 volumes of ice-cold 80/20 (v/v) methanol/water in 1.5 mL tubes by vortexing/trituration and then incubated for 20 minutes on ice. Samples were centrifuged for 10 minutes at 10,000 *g*, and the clarified supernatant was dried in a speed-vacuum/lyophilizer, and the dried pellet was stored at −80°C. Subsequent sample preparation for NMR spectroscopy was performed on the dried samples as described below.

For chlorophyll quantification in leaves, 10 days after germination (DAG) seedlings, grown as described above, were used. The glucanase activity assay was performed using crude extracts of 10 DAG leaves.

### Proton nuclear magnetic resonance spectroscopy

For the ^1^H NMR analysis, aforementioned extracted samples were resuspended to a final volume of 600 μl in D_2_O, with 0.35 mM sodium trimethylsilyl [2,2,3,3-d4] propionate (TSP), added to each lyophilized, titrated extract for chemical shift calibration. All sample preparations were performed over a period of two days and samples subsequently stored at 4°C. Quantitative ^1^H-NMR spectra were recorded at 800 MHz and 300 K on an Avance III spectrometer (Bruker Biospin, Wissembourg, France) using a 5-mm ATMA broadband inverse probe. One-dimensional ^1^H experiments, with a mild presaturation of water resonance, were performed with a 90° pulse angle.

The spectra were processed and analyzed with Chenomx NMR Suite 8.1 software (Chenomx Inc., 2014). Fourier-transformed spectra were multiplied with an exponential weighting function corresponding to a <u>line-broadening</u> of 0.5 Hz. All the spectra were manually phase-corrected, baseline optimized, and referenced to TSP. The metabolite peaks of the processed spectra were analyzed and assigned to their chemical shifts using the built-in Chenomx and the Human Metabolome Database (Wishart et al., 2018). The assigned metabolites were compared and confirmed through chemical shift values of other NMR based metabolomics studies performed in Arabidopsis (Hendrawati et al., 2006; Gromova and Roby, 2010; Augustijn et al., 2016), and through comparison with the Metabolomic Repository Bordeaux (MeRy-B) database (Deborde and Jacob, 2014). The concentrations of the assigned metabolites were determined with the Chenomx software and the concentration of the internal standard sodium trimethylsilylpropanesulfonate (DSS) as reference (Gromova and Roby, 2010).

### Statistical analysis of NMR spectroscopy datasets

Metabolite concentrations in the leaf and root extracts of the different experimental conditions were analyzed using a multivariate statistical analysis based on previously established methods (Krishnan et al., 2014; Khan et al., 2009). Briefly, a linear model fit was determined for each analyte using the LIMMA package in R (R Core Team, 2018) and lists of analytes with the most evident differential levels between the groups (control vs. treatment; leaves vs. roots; growth periods and ES7 concentration) were obtained. Significant analytes were selected by a two-step process. First, the initial data set consisted of measurements of all the analytes for which a signal was detected for at least one feature (e.g., control group at 4 DAG) for one condition. Second, the data from all the comparisons were combined into a single data set. The resulting combined data set consisted of analytes that exhibited modulation for at least one experimental comparison tested. Differential measurements within groups of samples, i.e. control samples at a particular day and ES7 treated samples, were detected by an *F* test. *P*-values for different analytes were transformed to compensate for multiple comparisons using the False Discovery Rate (FDR) adjustment for multiple comparisons using the Benjamini-Hochberg procedure (Benjamini and Hochberg, 1995; Benjamini et al., 2001). Fold changes were derived from multivariate statistical analysis. This analysis allowed a comparison of more than one statistical variable in one group, and therefore increases the statistical dimensionality of the data to provide a more meaningful value for fold change and adjusted p-values across multiple comparisons. The threshold for significance was a *p*-value < 0.1 for all tests with a fold change of (log2) > 1.5, unless otherwise stated in the specific analysis. All the analyses and plots were produced using a combination of Bioconductor and R (Gentleman et al., 2004; R Core Team, 2018).

### Spectrophotometric assays

Glucanase activity assays were performed according to Choudhury et al. (2010). Leaves of 10 DAG Arabidopsis seedlings harvested from agar plates were homogenized in liquid nitrogen with 50 mM sodium acetate buffer (pH5.2) containing 1 mM PMSF in a 1:1 w/v ratio using mortar and pestle. The homogenates were then filtered through Miracloth (MilliporeSigma, Burlington, MA, USA), and subsequently cleared by 1000*g* centrifugation at 4°C for 2 min. The clear upper phase of the lysate was desalted by size exclusion chromatography using a PD MiniTrap G-25 prepacked column (GE Healthcare, Chicago, IL, USA) with assay buffer as eluent. The protein content was measured by the Bradford protein assay (Bradford, 1976) and extracted proteins were used for the glucanase assay and background level estimation as described below. The assay mixture of 100 μL contained 50 μL desalted crude extract, 1 μL of DMSO or DMSO containing ES7, 19 μL of DI water, and 30 μL laminarin (TCI America, Portland, OR, USA) to yield a final concentration of 15 g/L as substrate. The standard curve was built with 3.125 to 100 μg of glucose dissolved in the assay buffer. Assays were performed at 50°C for 45 min, and terminated with 900 μL 3,5-dinitrosalicylic acid reagent at 85 °C for 10 min, before measuring the absorbance at 510 nm on a spectrophotometer (UV-1700, Shimadzu, Kyoto, Japan). Background levels of reduced sugars in the assay were determined using boiled protein extracts as reference. Leaf chlorophyll content was quantified according to established protocols (Porra et al., 1989). Briefly, weighed Arabidopsis leaves (20-40 mg) were extracted with 400 μL methanol/chloroform (2:1, v/v) for 1 h, and 300 μL water with 125 μL chloroform was added into the mixture to facilitate phase separation. After centrifugation at 10000*g* for 5 min, the lower chloroform phase was dried by air and resuspended in methanol. Chlorophyll a and b content was calculated from the sample absorbance at 665.2 nm, 652 nm, and 750 nm, with the extinction coefficient in methanol, using the formula indicated by Porra et al. (1989).

## Results

### Phenotypic responses of ES7 treated Arabidopsis seedlings

Arabidopsis seedlings were grown for 4, 5, 6, 10 days after germination (DAG) with 0 μM ES7 in the media as reference and control (**Fig 1A, B**). ES7 treatment was assessed by growing seedlings on media containing 3, 5, or 10 μM ES7 for up to 6 or 10 days (**Fig 1A, B**). The chosen ES7 concentration range was based on the previously established IC_50_ at 5 μM by root growth inhibition (Park et al., 2014). Given that ES7 reduces plant growth, we allowed the plants to grow for 6 or 10 days to ensure the availability of sufficient harvestable material for metabolite analysis. Seedlings were grown in vertical orientation in the plates to encourage directional root growth.

**Figure 1.**
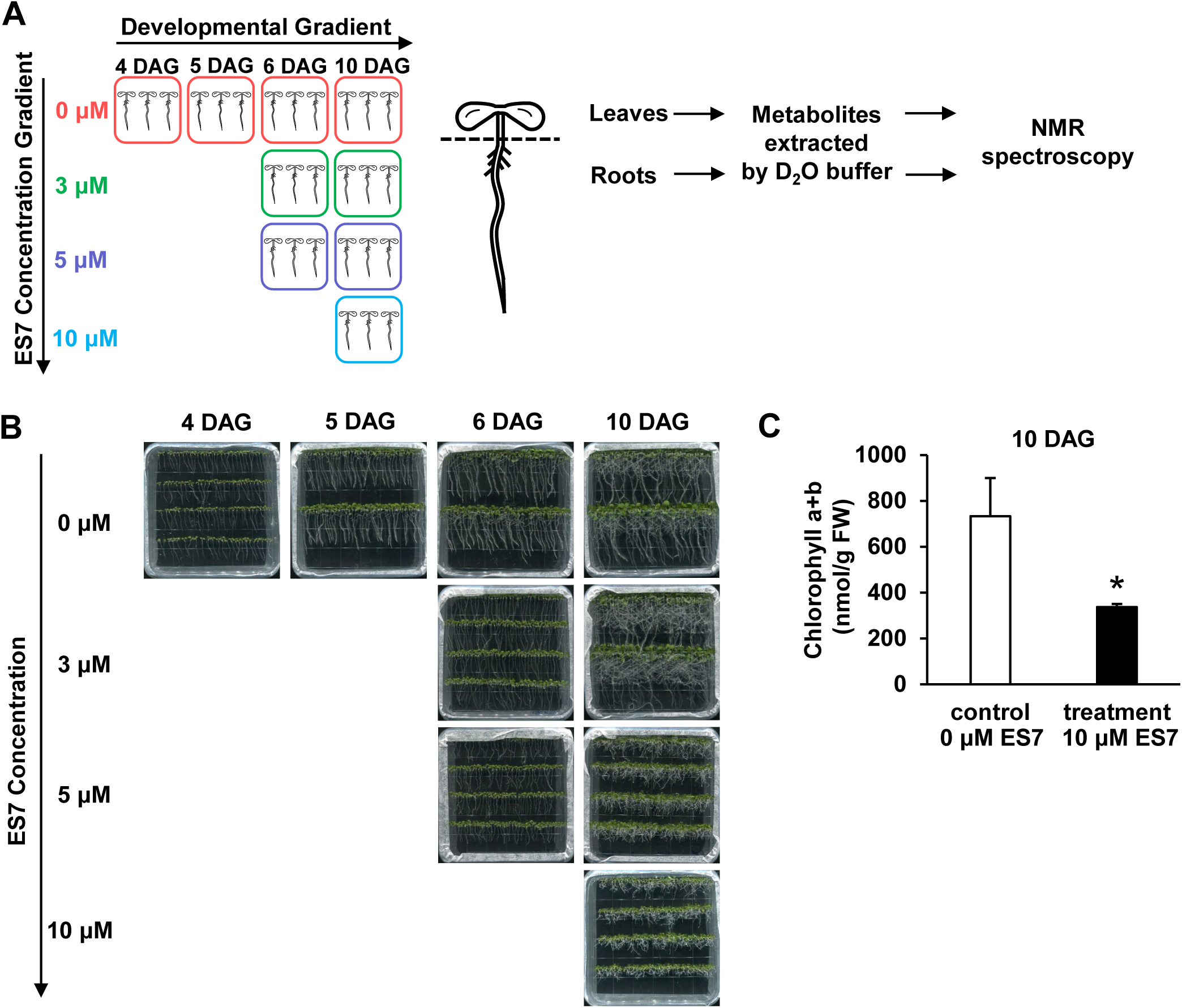
Experimental design and phenotypic responses of ES7 treatments. **(A) Experimental design of NMR Metabolomics analysis of ES7 effect on Arabidopsis**. Arabidopsis seedlings treated by 0, 3, 5, and 10 μM of ES7 and grown to 10 DAG, and 4, 5, 6, 10 DAG seedlings are collected as developmental control. Each graphic seedling represents one biological replicate of pooled seedlings from one plate. Leaves and roots are extracted by deuterated water, and subjected to NMR spectroscopy analysis. **(B) Visual phenotype of ES7 effect on Arabidopsis seedlings**. Photos are taken before metabolite extraction for NMR analysis. **(C) Chlorophyll content of ES7 treated 10 DAG leaves compared to control**. Data are presented by mean±SD (n = 4), and asterisk indicates significantly lower than control (*p* <0.05, t-test).

Metabolites were extracted from leaves and roots and NMR spectra were recorded. ES7 treated seedlings exhibited consistently shorter roots, in a concentration-dependent manner, compared to untreated controls (**Fig 1B**), corroborating earlier observations (Park et al. 2014). Notably, loss of gravitropism was observed in ES7 treated 10 DAG samples. The aerial part of the leaves was similarly affected, as indicated by its diminished growth. To assess the impact on the leaves, we measured the chlorophyll content of 10 DAG seedlings treated with 10 μM ES7. The leaf chlorophyll content showed a >50% reduction compared to the untreated control (**Fig 1C**), indicating a significant loss of photosynthetic activity.

### The effect of ES7 treatment is greater than the effect of developmental stages on the Arabidopsis metabolomes

Generally, a difference in the NMR spectra representing the metabolite profiles was observed for the different tissues, as shown by spectral excerpts of 4 DAG seedlings (**S1A Fig**). A partial least squares discriminant analysis (PLS-DA) of all the untreated leaf and root samples (**S1B Fig**), underscores the prominent difference between leaf and root metabolites. Biological triplicates of control metabolomes for each time point were tightly correlated with each other, demonstrating the robustness of our analysis (**S2 Fig**). In general, the NMR spectra of the leaves tend to have additional spectral features at the aromatic region and beyond (>7.00 ppm) in comparison with the NMR spectra of the roots (**S2 Fig**). Furthermore, in considering all the untreated samples (all DAGs, **Fig 2A**), the metabolites of the root samples cluster much tighter than that of leaves (**S1B Fig**), indicating a larger variation in the leaf samples. PLS-DA on the different DAG control metabolomes showed a difference across the development gradient (**Fig 2A**, PC1 = 16.8% and 18.8% for leaves and roots, respectively).

**Figure 2.**
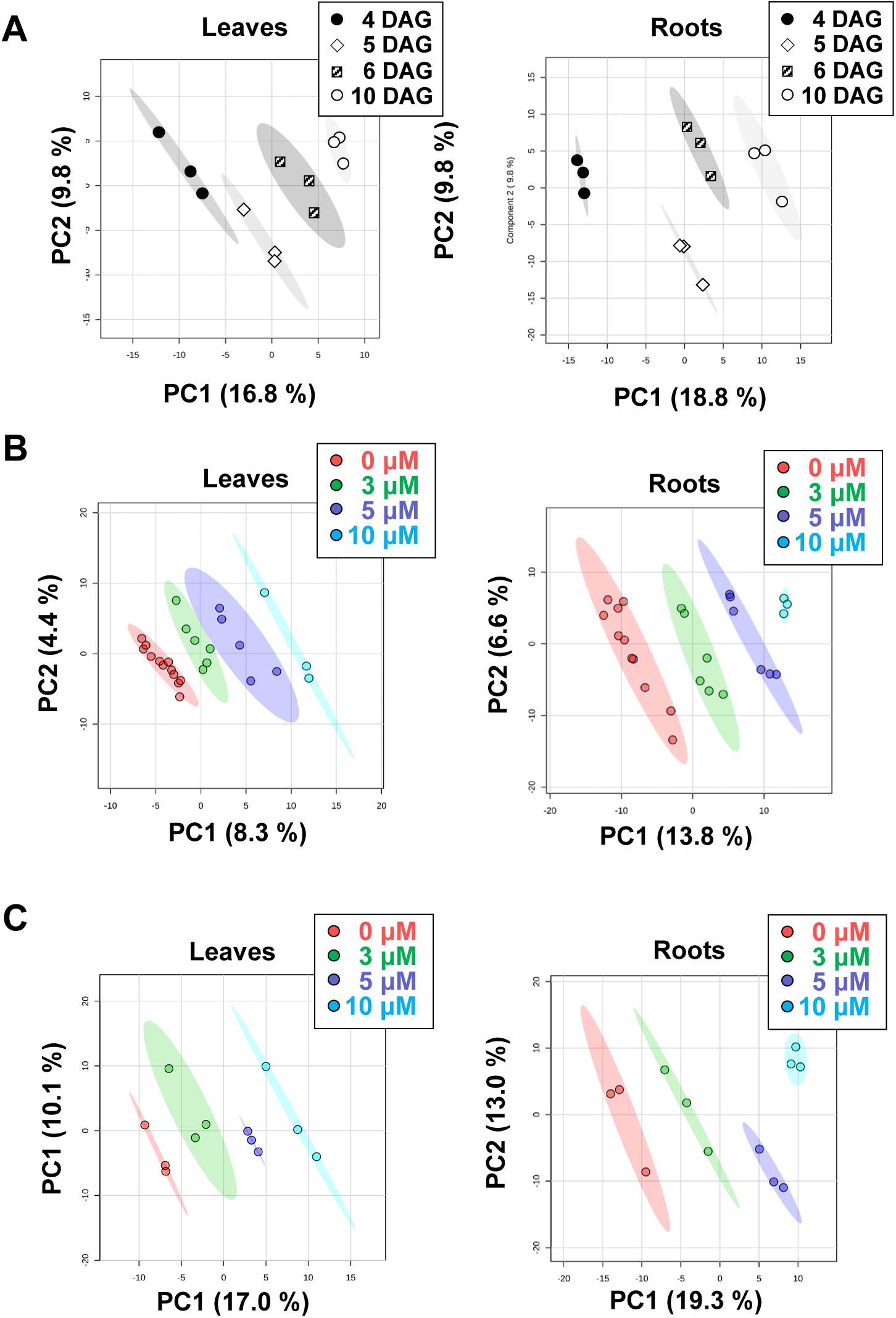
Classification of NMR based metabolomics of the ES7 treated seedlings versus control. **(A) PLS-DA analysis of metabolomics data of Arabidopsis seedlings at different DAG. (B) PLS-DA analysis of all NMR data of this experiment. (C) PLS-DA analysis of 10 DAG samples**. In **(B)** and **(C)** Each dot represents one biological replicate as indicated in **Fig 1A** with the same color scheme, and ellipses of 95% confidence region are over the classifications of biological replicates with respective ES7 concentrations.

Given that both plant development and chemical inhibition are contributing to the observed phenotypes, a series of statistical analyses were applied to determine if the ES7 treatment is the dominant factor contributing to the metabolite changes. We then analyzed the metabolite data of all 27 samples (**Fig 1A)** for roots and leaves, respectively, in order to assess the correlation of metabolomes across the developmental gradient and ES7 treatment. For both roots and leaves, PLS-DA of 0, 3, 5, and 10 μM ES7 entries are grouped tightly into areas of 95% confidence regions, marked by ellipses, and are well separated from the collective metabolome of the 12 untreated controls, across the developmental gradient (**Fig 2B**). Even at the lowest used ES7 concentration of 3 μM, no cluster overlap was observed with the control samples (**Fig 2B**, PC1 = 8.3% and 13.8% for leaves and roots, respectively). Focusing on the 10 DAG metabolomes (**Fig 2C**), with 0, 3, 5, and 10 μM ES7 concentrations, a clear separation of clusters was observed, accounting for the chemical treatment. This unequivocally demonstrates that the metabolic differences between samples are primarily explained by ES7 concentration rather than the developmental gradient of the sample.

### ES7 disrupts the homeostasis of primary metabolism in Arabidopsis

After verifying that ES7 treatment caused significant changes in metabolome composition, surpassing that of the developmental gradient, we focused on identifying the most prominently altered metabolites. Through PLS-DA analysis (**Fig 2**), the sub-groups of samples either due to developmental gradient or the concentration gradient of ES7 are discriminated. To increase the sensitivity of metabolite detection within the biomarker window and the statistical power of biological replicates, we focused on comparing ES7 effect between treated and untreated samples in the leaves or roots, independent of the developmental stage. Metabolite level changes were considered significant when the threshold of fold-change (log2) > 1.5 with corresponding adjusted p-values < 0.1. Fifty-three metabolites were significantly changed upon ES7 treatment in leaves or roots (*p* < 0.1), as listed in **S1 Table** (ES7 treatment n =15, control n =12). Individual compounds in **S1 Table** was first searched in the Kyoto Encyclopedia of Genes and Genomes (KEGG, genome.jp/kegg, (Kanehisa et al., 2002)) for their roles in Arabidopsis metabolic pathways. Their putative involvement and role in the biochemical pathways of leaves and roots are summarized in **Fig 3**, with a network map adapted from KEGG pathways. Detected metabolites can be categorized into components and derivatives of seven major metabolic pathways, including carbohydrate metabolism (Ruan, 2014), glycolysis and Krebs cycle (Plaxton, 1996; Sweetlove et al., 2010), glycerophospholipid metabolism, branched-chain amino acid metabolism (Binder, 2010), glycine, serine, and arginine metabolism (Bourguignon et al., 1998; Hildebrandt et al., 2015), shikimate pathway (Maeda and Dudareva, 2012), and pentose phosphate pathway (Kruger and Von Schaewen, 2003), as indicated in **S1 Table**. The average concentrations of the metabolites in the control or ES7 treatment are listed for leaves and roots, respectively (**S2 Table**), and individual biological replicates are shown in the boxplot to indicate the variation of each of the metabolites in the treatment (**Fig 4**). Most of the detected metabolites are considered to be part of primary metabolism pathways, involved in central plant growth and developmental processes.

**Figure 3.**
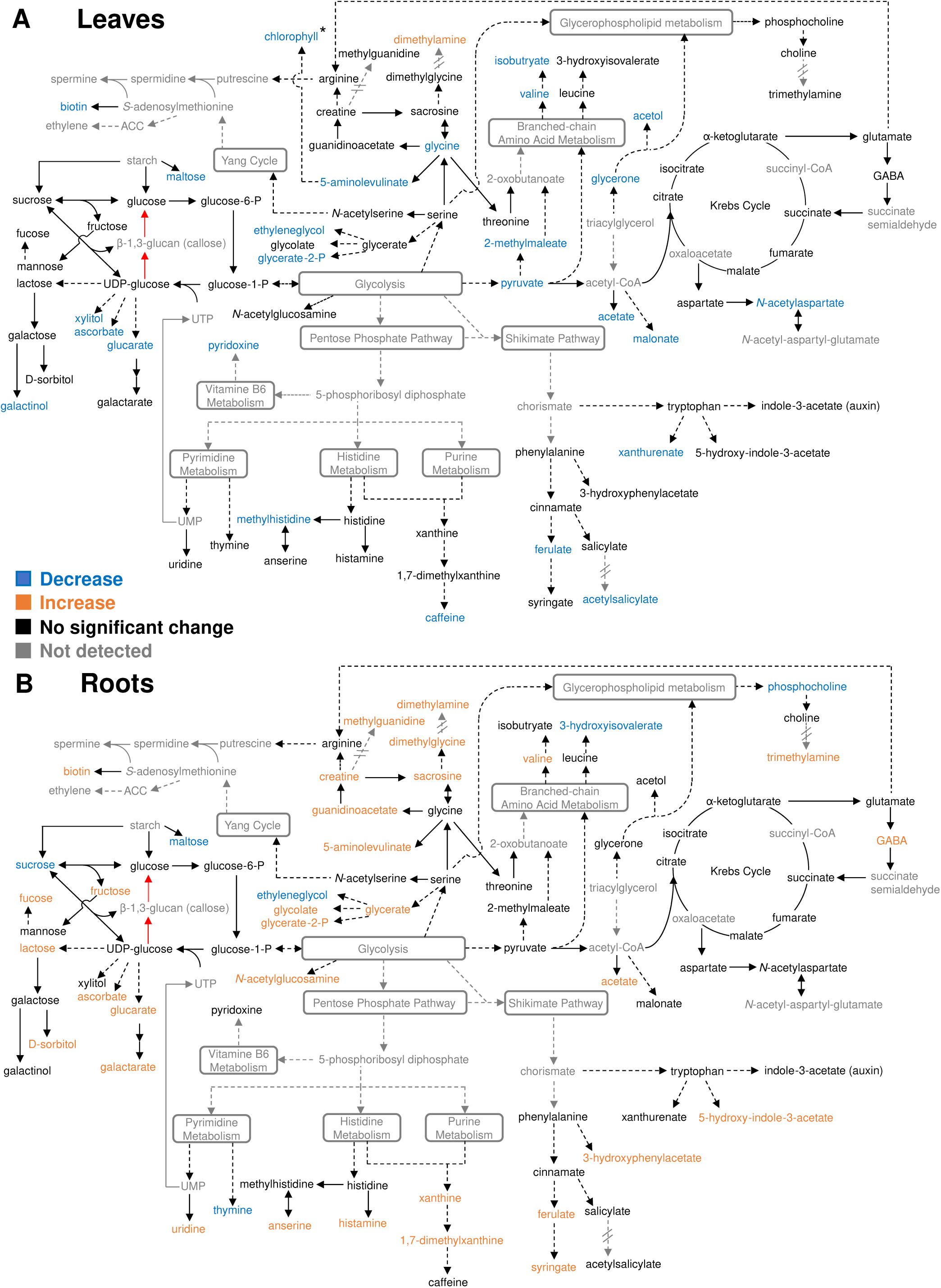
Metabolic pathway map showing metabolites significantly changed upon ES7 treatment in leaves. **(A) and roots (B) of Arabidopsis seedlings**. Backbone pathways are adapted from KEGG (genome.jp/kegg). Each solid arrow indicates one enzymatic step, and each dashed arrow indicates multiple enzymatic steps. Two red arrows represent callose synthase and β-1,3-glucanase respectively. Grey rounded rectangle indicates major pathways of multiple steps, and compounds in grey are not detected in the analysis. Dashed arrows with double cross lines indicate metabolic steps not reported in plants. Compounds in orange indicate significant increase, and ones in blue indicates significant decrease, and ones in black indicate no significant change upon ES7 treatment (significant as *p* < 0.1 in multivariate analysis, details indicated in **S1 Table**). *Chlorophyll change is inferred from the analysis of **Fig 1C**.

**Figure 4.**
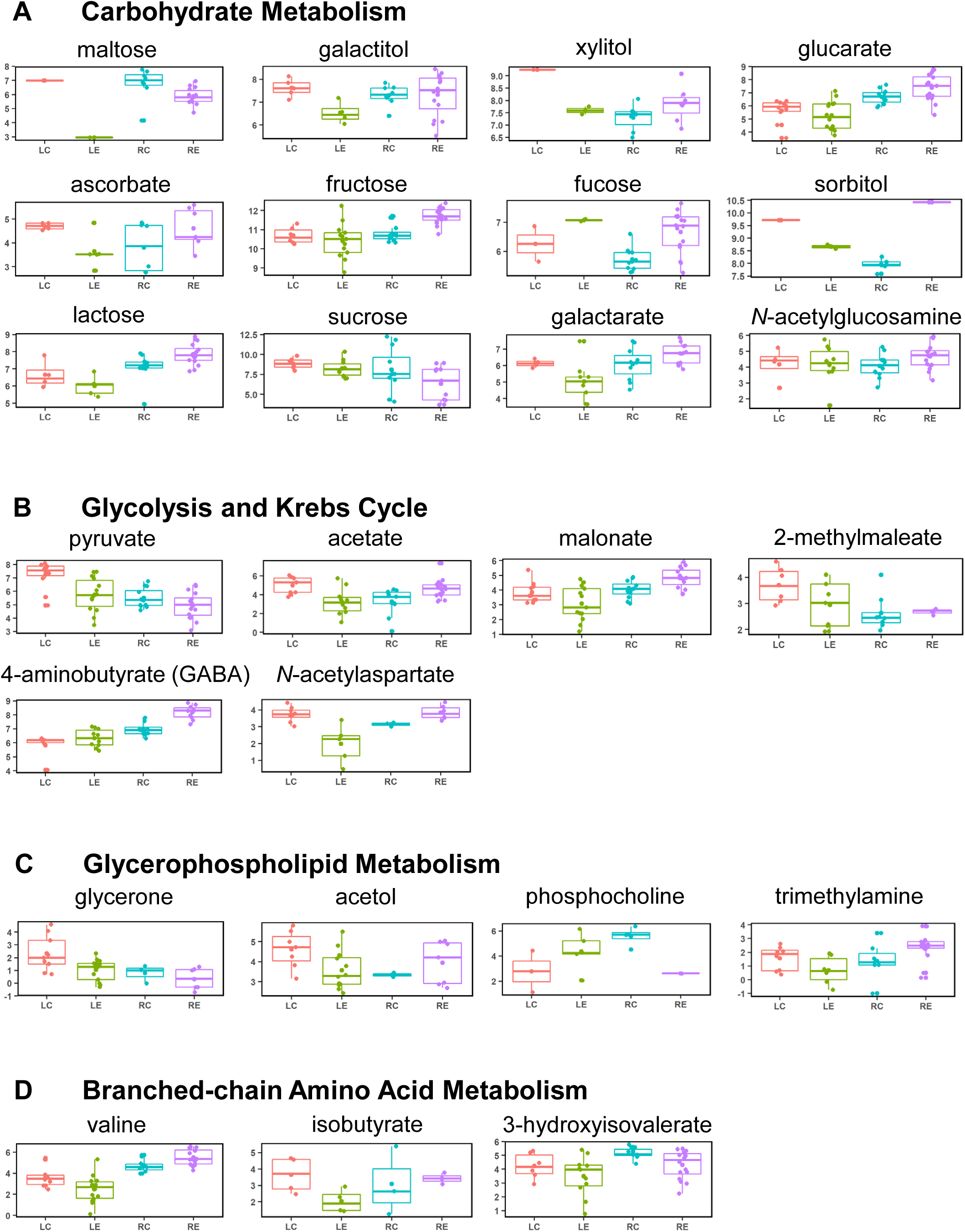
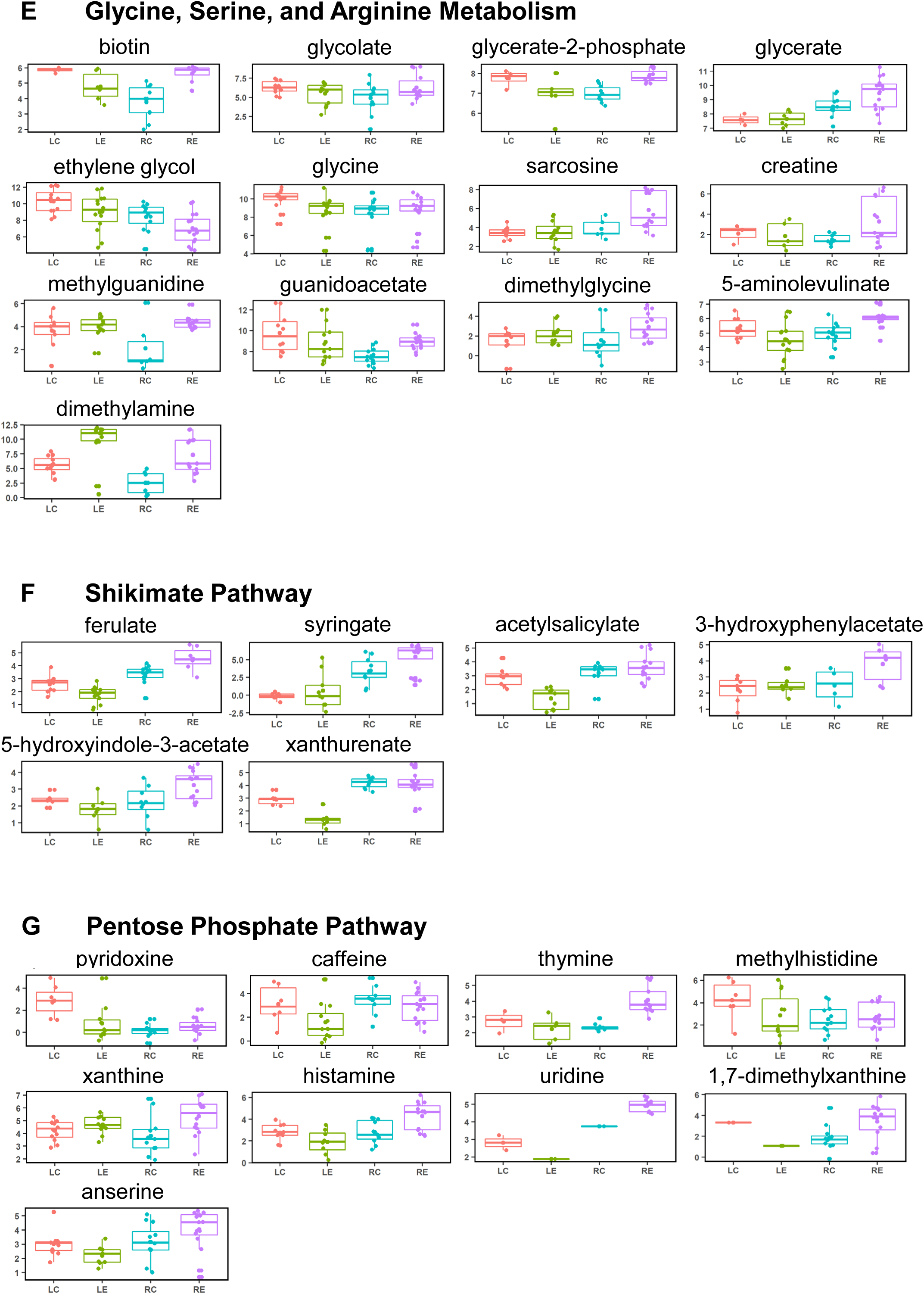
Differential levels of NMR estimated metabolites between the control and treated samples in roots and leaves. Metabolites are categorized into their major involved pathways or upstream pathways. Each panel lists a specific metabolite for LC (Control leaves, n = 12), LE (ES7 treated leaves, n = 15), RC (Control roots, n =12) and RE (ES7 treated root, n = 15). Concentrations are given in mM/g (log2) with respect to the internal standard DSS.

There were four metabolites with uncharacterized biosynthetic pathways: methylguanidine, dimethylamine, trimethylamine, and acetylsalicylate. These metabolites were plotted in the metabolite map based on structural similarity with putative precursors, as indicated by dashed lines with a break in the arrows (**Fig 3)**. Methylguanidine and trimethylamine have been independently reported in plant metabolomes using NMR spectroscopy (Wang et al., 2016) and liquid chromatography-tandem mass spectrometry (Tsukaya et al., 2015).

A dominant part of the modulated root metabolites showed an increase upon ES7 treatment (**Fig 3B**), in contrast to a decrease in leaves (**Fig 3A**). This difference likely reflects the different metabolic needs and compensatory mechanisms in aerial tissues and roots. The levels of most reduced sugars and their derivatives showed increased accumulation in roots and a reduction in leaves upon ES7 treatment, with the exception of maltose and sucrose showing a decrease in roots (**Fig 4A, S2 Table**). The physiological concentration of maltose has been shown to maintain membrane potential and protect the photosynthetic electron transport chain *in vitro* (Kaplan and Guy, 2004). The decrease of maltose in both roots and leaves upon ES7 treatment may be an indication of disrupted primary metabolism homeostasis. In roots, ES7 induced increases in compounds upstream of polyamine biosynthesis, including creatine, guanidinoacetate, and sarcosine (**Fig 4E**), and they are derivatives of glycine, serine, and arginine metabolism. ES7 treated roots also exhibited an increase in 4-aminobutyrate (GABA) (**Fig 4B**), a derivative of Krebs cycle, and its production is closely related to *in vivo* polyamine levels under stress (Zarei et al., 2016; Shelp et al., 2012). ES7 also induced changes of ferulate, syringate, 5-hydroxyindole-3-acetate, and xanthurenate in both roots and leaves (**Fig 4F**). As products of shikimate pathway, these compounds are phenylpropanoid and tryptophan derivatives related to the precursors for the biosynthesis of plant hormones, auxin and salicylic acid (Zhao, 2014; Dempsey et al., 2011). These metabolite changes suggest a pronounced effect of ES7 on the plant hormone biosynthesis pathways.

### The pool of UDP-glucose and glucose are not significantly affected by ES7

We did not detect significant changes in the direct metabolic substrate and degradation product of callose (β-1,3-glucan), namely UDP-glucose and glucose, upon ES7 treatment (**Fig 3**). This shows that inhibition of cytokinesis specific callose deposition does not cause a global change in the precursor pool. This is not surprising, given that UDP-glucose is involved in many metabolic activities and other cell wall polysaccharide biosynthesis steps, like starch and cellulose, beyond the transient accumulation of callose at the cell plate.

Callose deposition is regulated by the activity of both callose synthases and β-1,3-glucanases (**Fig 5A)** (Levy et al., 2007). Arabidopsis has twelve homologs of callose synthases (Verma and Hong, 2001) and fifty homologs of β-1,3-glucanases (Doxey et al., 2007). It is plausible that ES7 enhances the activity of cytokinetic β-1,3-glucanase(s), leading to higher phragmoplast callose degradation, thereby inhibiting polymer availability. To test this hypothesis, we examined the total glucanase activity modulation in Arabidopsis crude extracts following treatment with ES7. At 10 μM and 100 μM ES7, total glucanase activity in the crude extracts of Arabidopsis seedlings did not show a statistically significant change compared to the DMSO control (**Fig 5B**). This strongly suggests that global β-1,3-glucanase activity is not affected by ES7, and corroborates our NMR observations that ES7 does not cause a significant change in the direct metabolic substrate and degradation product of callose.

**Figure 5.**
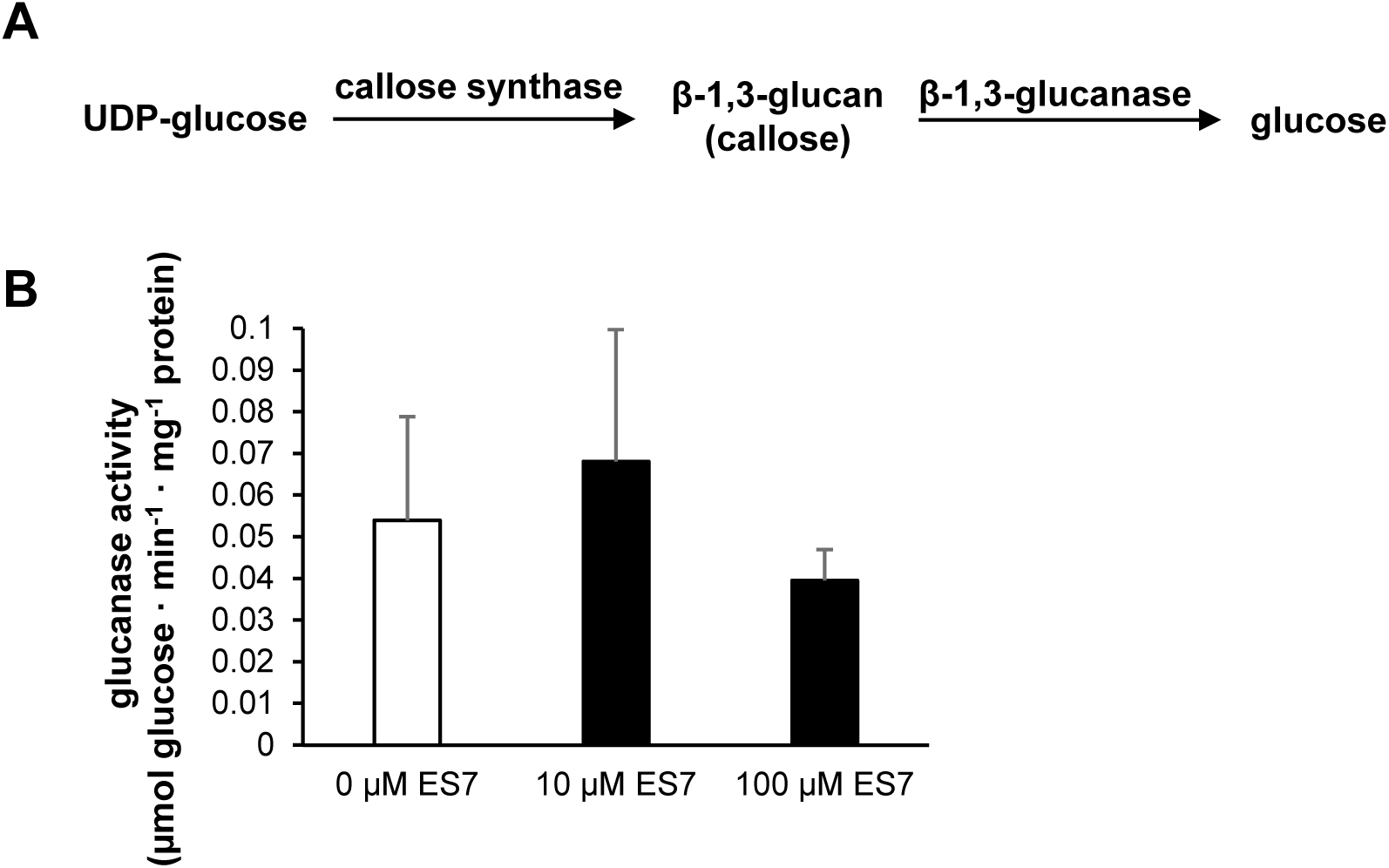
Total glucanase activity with or without the presence of ES7. (A) Callose synthase and β-1,3-glucanase regulate the dynamic equilibrium of the callose deposition in vivo. Park et al. (2014) demonstrated callose synthase activity is likely to be inhibited by ES7. **(B) Total glucanase activity in the crude extracts of Arabidopsis seedling leaves**. Activities are measured by the amount of glucose generated from incubating desalted crude extract with laminarin in the assay. DMSO, 10 μM ES7, or 100 μM ES7, are included in the assays respectively. Data are presented by mean±SD (n = 4).

## Discussion

### NMR-based metabolomics reveals small molecule induced changes

Plant metabolomics can be defined as the quantitative measurement of the time-related multiparametric metabolic response of the plants to environmental stimuli or genetic modification. The metabolomic information content is complementary to the genomic and proteomic approaches towards the interpretation of biological function and mechanism (Tian et al., 2007; Schauer and Fernie, 2006). Transcriptome and proteome analyses have been extensively recruited to study how tissue-specific responses are coordinated during growth, toward identifying mechanisms of plant development (Schad et al., 2005; Plant et al., 2012; Sreenivasulu et al., 2004; Müller et al., 2010; Ghatak et al., 2016). Especially in the field of gene expression analysis, recent analytical and methodical developments have allowed single-cell level-based analysis, which can help us understand how gene networks are organized and regulated at the cellular level (Nakazono et al., 2003; Brandt, 2002; Karrer et al., 1995). In the field of metabolomics, attempts to describe the metabolome of single cells have been based on tandem mass spectrometry techniques (Misra et al., 2014). In pioneering work, improving spatial resolution, a survey of the subcellular distribution of metabolites in the *Arabidopsis thaliana* leaf was performed towards cytosol, vacuole, and plastid fractions by Krueger and colleagues using GC-TOF/MS and LC/MS. The results provided a topological metabolite map, highlighting an initial step to analyze the metabolic dynamics between subcellular compartments (Krueger et al., 2011). While a metabolomic analysis can be performed using several mass spectrometry-based analytical chemistry techniques, NMR spectroscopy-based metabolomics analysis has some distinct advantages (Wolfender et al., 2019; Emwas et al., 2019). These may include easy and rapid sample preparation, free of derivatization analysis that offers high-throughput and quantitative analysis using NMR with a single internal standard (Izquierdo-García et al., 2011; Marti et al., 2013; Nagana Gowda and Raftery, 2017). In the field of plant biology, NMR techniques have been utilized in cell wall polymer characterization (Dick-Perez et al., 2012; Phyo et al., 2017; Wang et al., 2013; Yuan et al., 2016); however, the application of NMR-based metabolomics in plants is limited (Kim et al., 2011a), partially due to the insufficient availability of open-access NMR spectral libraries for plant metabolites (Ludwig et al., 2012; Johnson and Lange, 2015). External chemical stimuli NMR studies without tissue specificity have been conducted to evaluate the response of the plant defense inducer benzothiadiazole (Hien Dao et al., 2009), the hormone methyl jasmonate (Hendrawati et al., 2006), the trafficking inhibitor Sortin 1 (Orr et al., 2014), and the pathogen *Verticillium dahliae* (Su et al., 2018). The Arabidopsis metabolic profiles, with and without pharmacological inhibition, presented here can contribute to an enhanced understanding of general plant responses to small molecule stimuli.

### Tissue-specific metabolomes to dissect small molecule induced responses in plants

We combined tissue-specific metabolite changes as a response to a small molecule, Endosidin 7, through NMR spectroscopy. In this study, ^1^H NMR spectroscopy profiling was applied to Arabidopsis leaves and roots separately, clearly demonstrating distinct tissue-specific differences quantified via PLS-DA (**S1 Fig**). Tissue-specific metabolomics has been utilized to study plant metabolism upon abiotic or biotic stress (Kim et al., 2011b; Ullah et al., 2017). Root and leaf metabolic responses towards sublethal cadmium exposure, high salt stress, and low potassium stress, respectively, were characterized via GC or LC-MS, and principal component analysis (PCA) of the three independent metabolomics studies consistently indicated differential responses in root and leaf tissue of Arabidopsis and Barley (Keunen et al., 2016; Zeng et al., 2018; Wu et al., 2013). In addition, Novák et al. (2012) studied the tissue-specific Arabidopsis auxin metabolome by LC-MRM-MS, for both roots and leaves of wild type and auxin over-producing lines. PCA analysis also indicated overproduction of auxin led to distinct modulation profiles in which up-regulation or down-regulation of the metabolites in leaves and roots did not follow a parallel pattern (Novák et al., 2012). In melons, tissue-specific metabolite changes via NMR were observed under the exposure to the powdery mildew disease, in mold-resistant rootstocks (roots) and susceptible watermelon scions (aerial parts), with both showing opposite changes in the level of root and leaf metabolites. The authors put forward the hypothesis that translocation of metabolites between rootstocks and scions through the vascular system is responsible for the antiparallel metabolome modulation (Mahmud et al., 2015). Our observed trend of metabolite level changes in leaves and roots upon ES7 treatment is also strikingly different (**Fig 3, S1 Table**), although the cellular phenotype of late cytokinesis arrest is observed in both leaves and roots after ES7 treatment (Park et al., 2014).

While one possible explanation of this antiparallel response in roots and leaves results from aberrant translocation of metabolites between plant organs under ES7 treatment, a second possible explanation is that intrinsic tissue-specific regulation of primary metabolic pathways is not concomitant in plant organs under stressed conditions.

The cited studies above, together with our presented data, indicate the importance of tissue-specific investigation to assess the coordinated responses of plant organs via NMR metabolomics.

### Long term ES7 treatment may disrupt Arabidopsis hormonal homeostasis

We performed a long-term ES7 treatment of 6 or 10 DAG in order to investigate the metabolic phenotype that captures both the long-term growth inhibition and the cellular phenotype of arrested cell division. Both the morphological phenotype and metabolite changes induced by ES7 treatment suggested a disruption of hormonal homeostasis. Loss of gravitropism is a sign of possible disruption of hormonal regulation, as gravitropism is regulated by crosstalk between auxin and other hormones (Nziengui et al., 2018; Philosoph-Hadas et al., 2005). We did not observe a significant change in indole-3-acetate (auxin) levels upon ES7 treatment; however, 5-hydroxyindole-3-acetate, a metabolite also putatively derived from tryptophan (Mahmud et al., 2015), showed a significant increase in roots (**Fig 4F**). Downstream of purine metabolism, cytokinin synthesized from the deoxyxylulose pathway (Zrenner et al., 2006; Mok and Mok, 2001) are likely to be affected upon ES7 treatment, implied by the increase of xanthine and 1,7-dimethylxanthine in roots (**Fig 3, Fig 4G**). We did not detect a significant change for the plant hormone salicylate, which is involved in both abiotic and biotic stress (Loake and Grant, 2007; Khan et al., 2015), but acetylsalicylate showed an over 60% reduction upon ES7 treatment in leaves (**Fig 4F, S2 Table**). Further, components of polyamine biosynthesis pathways are affected upon ES7 treatment. Altogether, our data suggest that the imbalance of certain hormone levels during prolonged treatment of ES7 could have induced the global changes of metabolite levels. Both auxin and cytokinin are responsible for the initiation of the G1/S phase transition prior to cell division, a prerequisite for cell division through the regulation of cyclin-dependent kinases (Wang and Ruan, 2013; Schaller et al., 2014; Hartig and Beck, 2006; Moubayidin et al., 2009). Also, auxin and cytokinin regulate the biosynthesis of each other via the direct regulation of biosynthesis genes and transporters (Rosquete et al., 2012). ES7-induced metabolome changes could represent a compensatory mechanism counteracting the inhibition of cytokinesis and plant growth, resulting from the imbalance of cytokinin and auxin.

This study also reveals that long-term treatment of ES7 disrupts the homeostasis of primary metabolism in Arabidopsis seedlings, likely via alteration of hormonal regulation. Microarray analysis revealed that 35 primary metabolism genes involved in light signaling, nutrient uptake, and photosynthesis are altered in Arabidopsis shoots treated by 5 μM isopentenyladenine (a synthetic cytokinin) for 4 days (Brenner et al., 2012; Che et al., 2002). In another long-term cytokinin treatment experiment, lettuce treated by benzylaminopurine or meta-topolin for 13 days showed reduced accumulation of photosynthetic pigment and inhibition of photosystem II activity (Prokopová et al., 2010). The phenotypes of ES7 treated seedlings, including a plethora of metabolite changes in primary metabolism (**Fig 3**) accompanied by loss of chlorophyll (**Fig 1C**), could also be explained through aberrant hormonal regulation, as chlorophyll synthesis is tightly regulated by the balance of auxin and cytokinin (Hudson et al., 2011; Liu et al., 2017).

Another possible long-term treatment effect of ES7 might be the disruption of lignin biosynthesis, as levels of ferulate and syringate, precursors of lignin biosynthesis, are affected upon ES7 treatment (**Fig 3** and **Table 1**). Aberrant lignin biosynthesis may further affect xylem development in Arabidopsis seedlings (Taylor-Teeples et al., 2015), interfering with nutrient translocation and potentially giving rise to opposite modulations of root and leaf metabolites.

Park et al. (2014) observed the ES7 inhibition effect on callose deposition at the division plate as short as two hours after pulse treatment. This time frame is in line with characteristic hormonal responses, in that gene expression or metabolite level changes are usually observed after a few hours of cytokinin or auxin induction (Talbott and Ray, 1992; Zürcher et al., 2013; Kull et al., 1978; Petit-Paly et al., 1999). Given that the cellular phenotype of cell plate disruption is observed as early as two hours when treated with a higher ES7 concentration of 50 μM, these conditions could be used in future NMR metabolomics studies to dissect the long-term effects from the short-term metabolite effects.

### Summary and perspectives

We performed a tissue-specific NMR-based metabolomics study of Arabidopsis leaves and roots at different developmental stages and treatment with various concentrations of the specific cytokinesis inhibitor, ES7. Metabolome analyses indicated that cytokinesis inhibition of ES7 likely also disrupts the homeostasis of primary metabolism and hormonal regulation. Our tissue-specific analysis highlighted the importance of spatial resolution in metabolite analysis. Given the complex interactions between metabolic pathways, future studies that allow higher spatial and temporal resolutions are important to unmask the different layers of interaction, particularly after exogenous stimuli. There are two ways to envision how this complex task can be addressed through future developments: a) automation through partially robotic extraction of the required substantial amounts of tissue, and b) improving metabolite identification and metabolic flux analysis using stable isotopic enriched and multidimensional NMR metabolomics with higher sensitivity leading to smaller required sample volumes (Deborde et al., 2017; Lane and Fan, 2017; Markley et al., 2017; Emwas et al., 2019). Such technological advances, in combination with cell synchronization, can uncover delicate metabolite changes during different cellular processes including cytokinesis. This study provides metabolomics references for early stages of Arabidopsis development, indicates multiple metabolic pathways affected by ES7, and highlights the relevance of tissue-specific investigation in plants for an accurate and comprehensive assessment of plant metabolome modulations upon small molecule stimuli.

## Acknowledgments

We thank Dr. Dan Kliebenstein for the discussion on the NMR metabolomics data. We thank Destiny Jade Davis for critical reading and editing of the manuscript. The work was supported by

U.S. National Institutes of Health grant SC3-GM125546 to V.V.K., U.S. National Science Foundation MCB-1818219 award to G.D., and U.S. Department of Agriculture award CA-D-PLS-2132-H to G.D.

## Supplemental Data

**S1 Figure.**
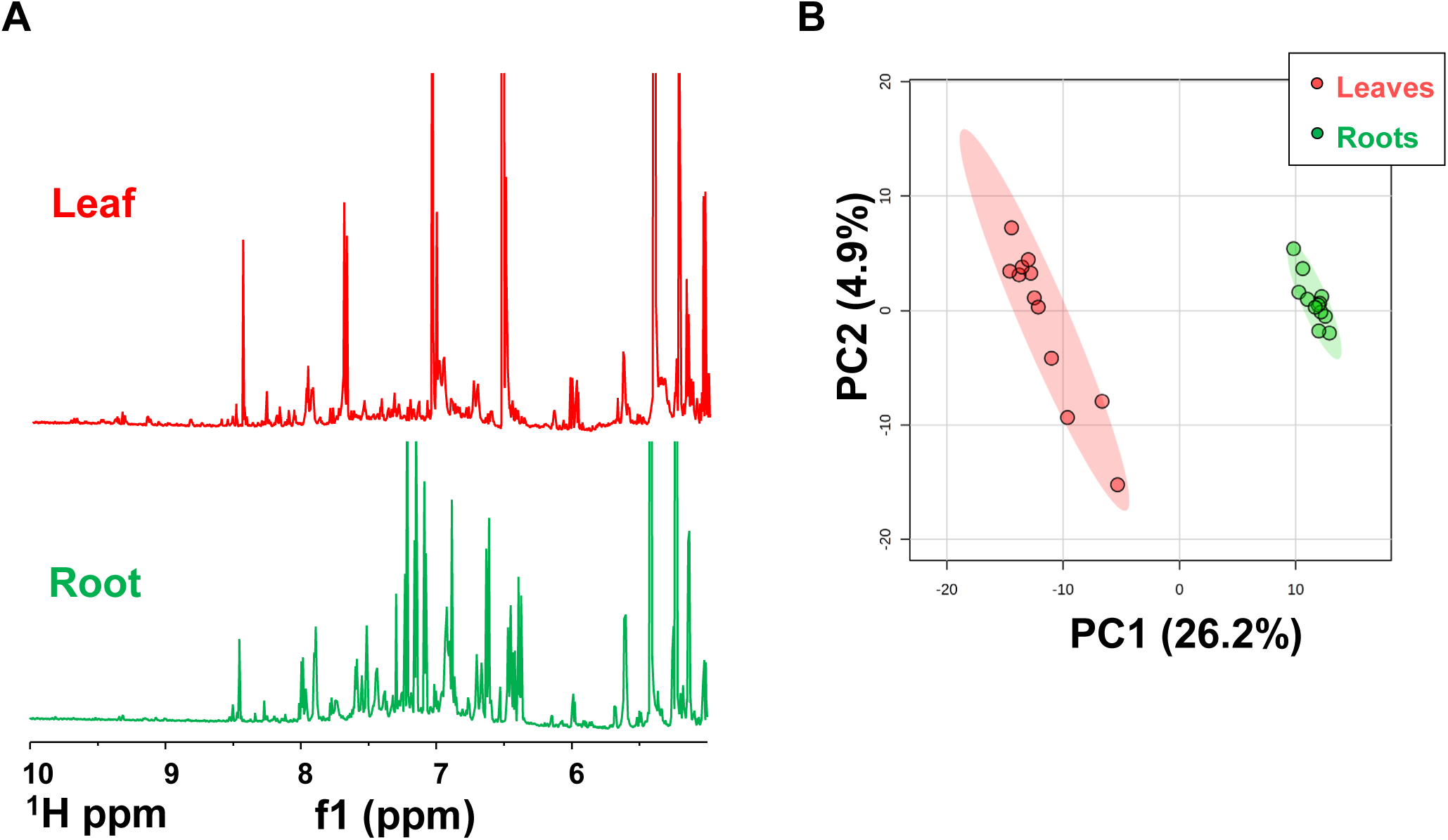
Characteristic differences between the NMR spectra of leaf and root samples. **(A) Low-field (> 5**.**00 pm) 1H NMR spectra of control samples on 4 DAG between the leaves (red) and roots (green). (B) PLS-DA analysis of leaves and roots metabolome**. All the biological replicates are included in the PLS-DA assay, and ellipses represent 95% confidence region of the classification.

**S2 Figure.**
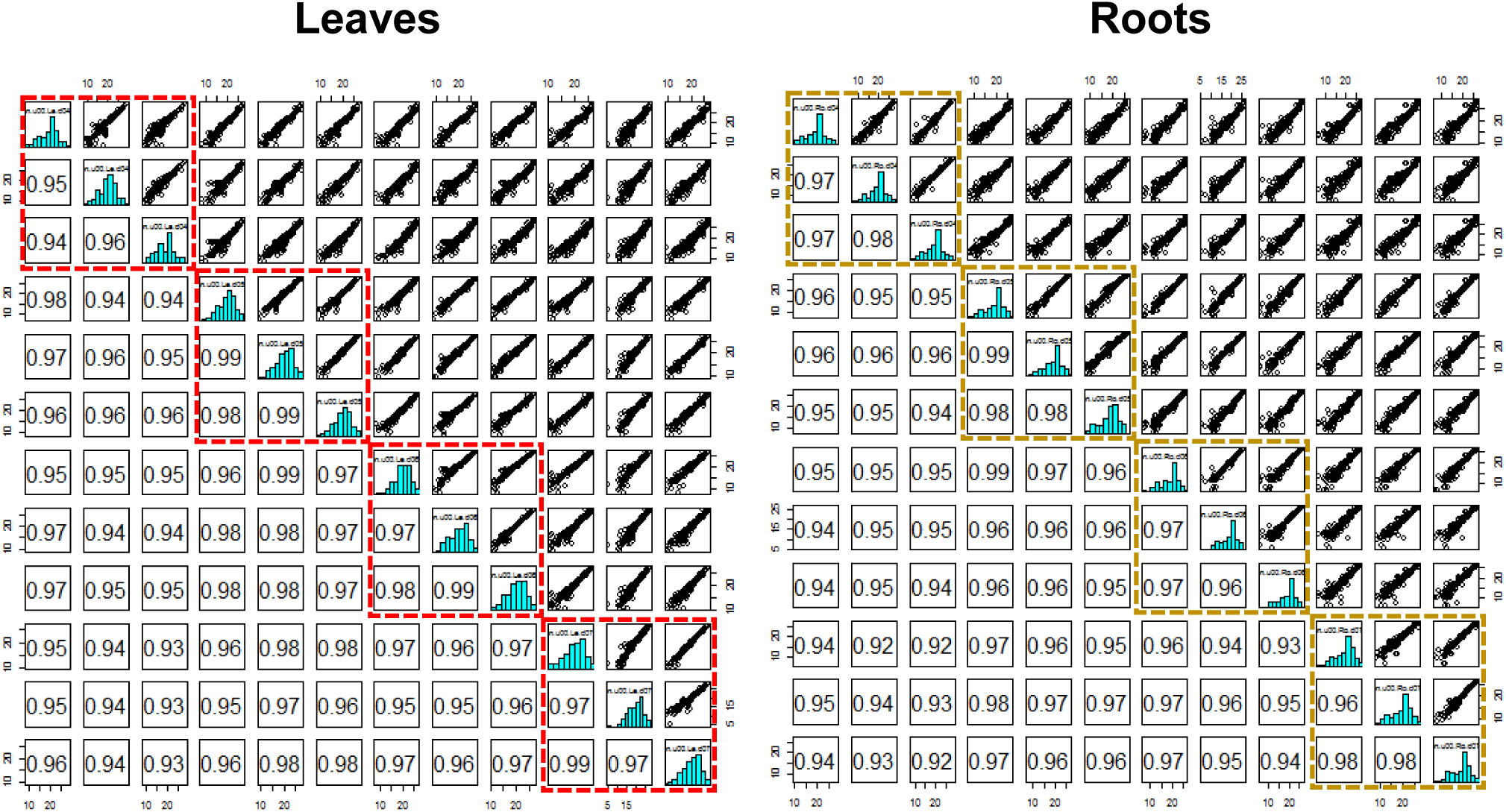
Correlation analysis among 4, 5, 6, and 10 DAG samples without ES7 treatment. Triplicates of each DAG are ranked with highest correlation with each other, highlighted in dashed square.

**S1 Table.**
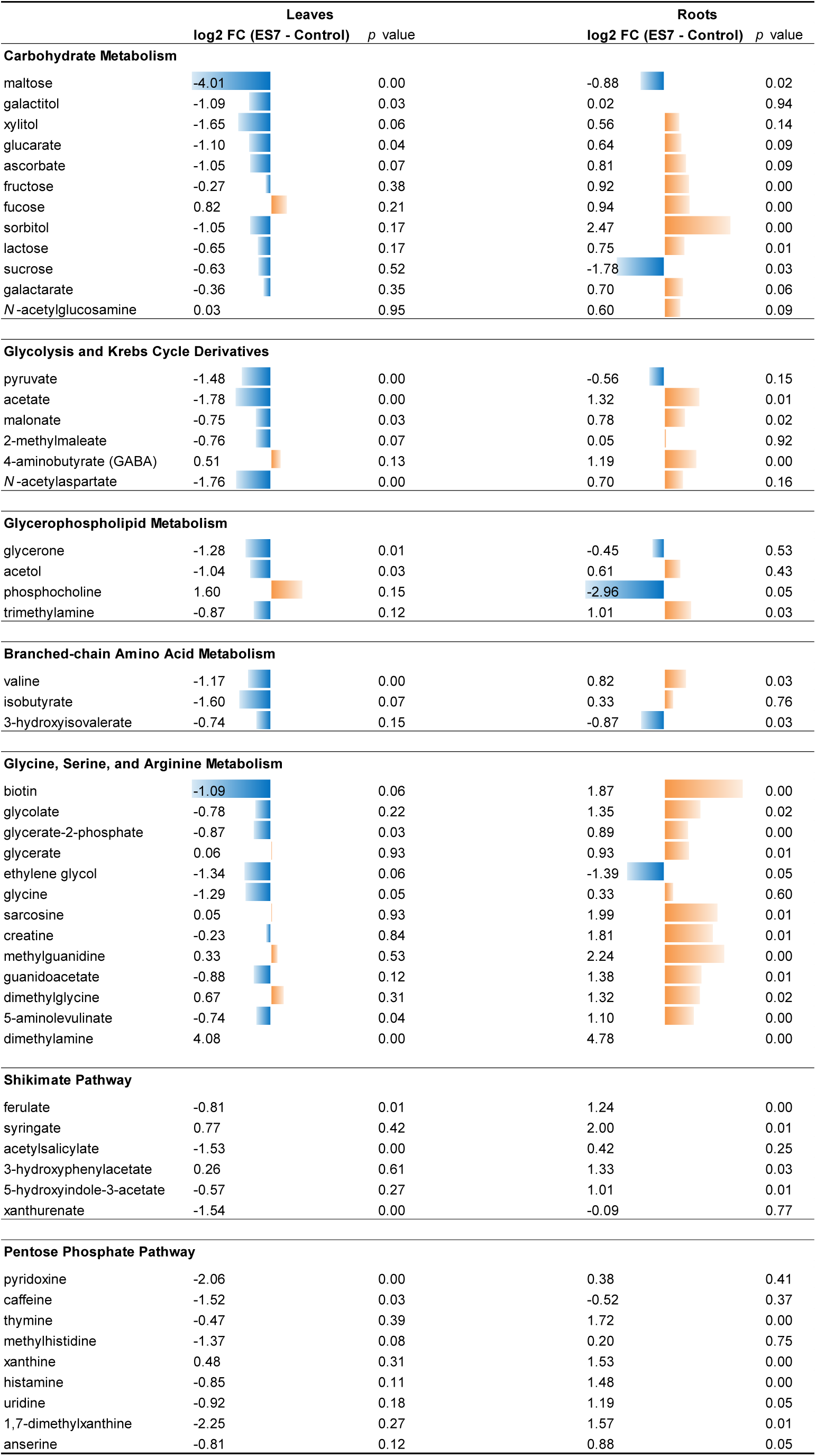
Metabolites significantly changed upon ES7 treatment in leaves and roots of Arabidopsis seedlings. Difference of metabolite levels between ES7 treatment (n=15) and control (n=12) are expressed by log2 fold change. Length of colored bar is proportional to the value, with negative values in blue, and positive values in orange, and threshold of 1.5 (log2) is used in multivariate analysis to calculate the *p* value of significance.

**S2 Table.**
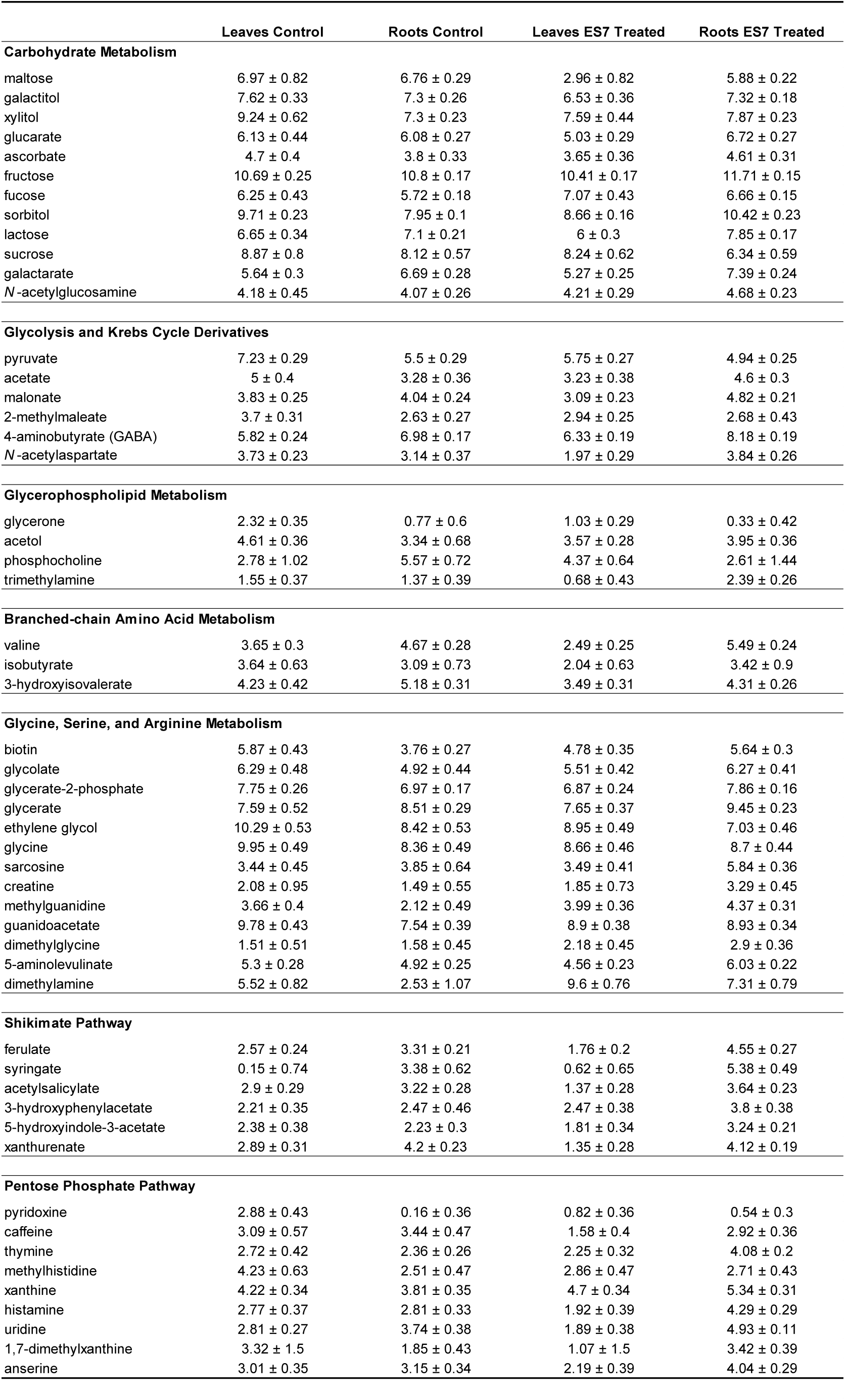
Quantification of metabolite levels changes upon ES7 treatment in leaves and roots. Concentrations (mean±SD) are given in mM/g (log2) with respect the internal standard DSS (n = 12 for controls, n = 15 for ES7 treated).

